# Auditory cortical generators of the Frequency Following Response are modulated by intermodal attention

**DOI:** 10.1101/633834

**Authors:** Thomas Hartmann, Nathan Weisz

**Affiliations:** Center of Cognitive Neuroscience, University of Salzburg, Salzburg, Austria; Department of Psychology, University of Salzburg, Salzburg, Austria

**Author notes:** correspondence: Dr. Thomas Hartmann, Paris-Lodron Universität Salzburg, Division of Physiological Psychology, Hellbrunnerstraße 34, 5020 Salzburg, Austria, Fax: +43 (0) 662 / 8044 - 5126.

**Keywords:** FFR, brainstem, auditory cortex, attention, MEG

## Abstract

The vast efferent connectivity of the auditory system suggests that subcortical (thalamic and brainstem) auditory regions should also be sensitive to top-down processes such as selective attention. In electrophysiology, the Frequency Following Response (FFR) to simple speech stimuli has been used extensively to study these subcortical areas. Despite being seemingly straight-forward in addressing the issue of attentional modulations of subcortical regions by means of the FFR, the existing results are highly inconsistent. Moreover, the notion that the FFR exclusively represents subcortical generators has been recently challenged. By applying these techniques to data recorded from 102 magnetoencephalography (MEG) magnetometers in 34 participants during a cross-modal attention task, we aimed to gain a more differentiated perspective on how the generators of the FFR are modulated by either attending to the visual or auditory input. In a first step our results confirm the strong contribution of also cortical regions to the FFR. Interestingly, of all regions exhibiting a measurable FFR response, only the right primary auditory cortex was significantly affected by intermodal attention. By showing a clear cortical contribution to the attentional FFR effect, our work significantly extends previous reports that focus on surface level recordings only. It underlines the importance of making a greater effort to disentangle the different contributing sources of the FFR and serves as a clear precaution of simplistically interpreting the FFR as brainstem response.

---

Analogous to other sensory modalities, neural activity in the auditory system is modulated by selective attention (Fritz et al. 2007; Frey et al. 2014; Mazaheri et al. 2014; Weise et al. 2016; Salo et al. 2017). Electrophysiological research has also focussed on oscillatory activity in cortical brain regions, revealing modulations in the auditory cortex similar to those reported in the visual domain (Händel et al. 2011) or somatosensory regions (Haegens et al. 2011). These findings point to alterations of gain in sensory cortical regions to select or ignore features respectively (e.g. gating by inhibition (Jensen and Mazaheri 2010)) that are modality-independent (Lee et al. 2012; Choi et al. 2013; Frey et al. 2014). Despite these similarities at the cortical level, compared to the visual modality the auditory system is characterized by a more extensive and complex subcortical architecture, including abundant efferent neural connections (Winer 2006; Suga 2008; Chandrasekaran and Kraus 2010; Terreros and Delano 2015). Within this efferent system, the primary auditory cortex is a hub region with direct efferent connections to all major subcortical areas (Winer 2006; Suga 2008; Chandrasekaran and Kraus 2010; Terreros and Delano 2015). In principle, auditory cortical processes could affect cochlear activity via only two synapses (Winer 2006; Suga 2008; Dragicevic et al. 2015). These corticofugal modulations are essential in adapting responses of subcortical neurons, for example, by modulating their spectral tuning curves (Suga 2008; Felix et al. 2018). However, the extent to which subcortical auditory brain regions are implicated in selective attentional modulation is not well established.

Recently, Slee and David (Slee and David 2015) reported attentional modulation of receptive fields of inferior colliculus (IC) neurons in ferrets. In humans functional magnetic resonance imaging (fMRI) has shown modulation of IC activity by selective auditory attention (Rinne et al. 2008; Riecke et al. 2018) and increases of BOLD activity with attentional demand in brainstem structures in an audiovisual attention task (Raizada and Poldrack 2007). Since all efferent connections to the cochlea are mediated via the Superior Olive, further suggestive support for subcortical attentional modulations can be derived from studies on otoacoustics emissions (OAEs), a proxy for outer hair cell activity in the cochlea. Attentional modulations of OAEs have been found when either the left or the right ear had to be attended (Giard et al. 1994), one out of two frequencies was task relevant (Maison et al. 2001) or attention had to be focused on the visual or auditory modality (Wittekindt et al. 2014). While limited in number, these studies in animals and humans suggest the sensitivity of brainstem structures to selective attention. However, the picture remains incomplete. Studies in the animal model are only suggestive that similar processes also exist in humans. At the same time invasive recordings from brainstem structures are not feasible in healthy humans. Studies using fMRI studies are non-invasive and provide excellent spatial resolution. Yet, scanner noise creates a challenging environment for such studies, and the technique is also not well suited for some populations in which the study of brainstem processes may be of interest (e.g. cochlear implant patients). Furthermore, the aforementioned complex auditory corticofugal architecture strongly suggests complex interactions of the areas and nuclei involved which can only be captured with recording methods providing high temporal resolution. Therefore other methods are needed to complement the invasive and neuroimaging approaches.

A popular method to noninvasively assess auditory neural activity with high temporal resolution is to use magnetoencephalography (MEG) and/or electroencephalography (EEG). Convincingly capturing attentional modulations of subcortical auditory regions using these techniques has proven challenging, however. Many initial studies focussed on the components of the auditory brainstem response (ABR) (Jewett et al. 1970). The ABR is the evoked response to a high number of repetitions (typically >5000) of a short sound like a click. The ABR peaks have been related to the processing of the sound at the various subcortical nuclei (Jewett et al. 1970; Don and Eggermont 1978; Møller et al. 1981; Møller and Jannetta 1983; Boston and Møller 1985; Møller and Burgess 1986; Chandrasekaran and Kraus 2010; Skoe and Kraus 2010). It has been used to study plasticity processes of the subcortical auditory system (Musacchia et al. 2007; Tzounopoulos and Kraus 2009; Chandrasekaran and Kraus 2010; Chandrasekaran et al. 2014; Kraus and Nicol 2014) and is widely applied in clinical settings (Eggermont et al. 1980; Møller and Møller 1983; Eggermont and Don 1986; Eggermont and Salamy 1988; van Straaten 1999; Stipdonk et al. 2016). However, attentional modulation of ABR components have not been found thus far (Woldorff et al. 1987; Hackley et al. 1990).

Another way to use EEG to eavesdrop on subcortical activity is to investigate neural responses to complex sounds such as simple consonant-vowel combinations (e.g. /da/ sound). While the transient part is equivalent to the classical ABR, the sustained part is an oscillatory response that is strictly phase-locked to the stimulus, in particular its fundamental frequency (F0) (Greenberg 1980; Galbraith et al. 1995; Russo et al. 2004; Akhoun et al. 2008; Chandrasekaran and Kraus 2010; Skoe and Kraus 2010). This sustained response, commonly named the frequency-following response (FFR), is assumed to be mainly generated by subcortical auditory nuclei, with the IC playing a central role (Worden and Marsh 1968; Batra et al. 1986; Chandrasekaran and Kraus 2010). Support for this notion has come from studies in animals (Marsh et al. 1974; Rouiller et al. 1979; Liu et al. 2006; Wallace et al. 2007). Given these findings and the general notion that the auditory cortex does not track or hardly tracks frequencies beyond 100Hz (Kuwada et al. 2002; Chandrasekaran and Kraus 2010), the FFR has thus been seen as a proxy to subcortical auditory activity. Past studies on the attentional modulation of the FFR have found inconsistent results. While some find in favor ((Galbraith et al. 2003; Hoormann et al. 2004; Lehmann and Schönwiesner 2014); for an alternative innovative method exploiting the neural response to the F0 showing attentional modulations, see (Forte et al. 2017)), negative results have been reported as well (Varghese et al. 2015). It should be noted that all mentioned studies used data from a small number of EEG electrodes, sometimes only one. Since the subcortical generation of the FFR has been widely accepted, the presence or absence of attentional modulations in the aforementioned studies has been attributed to subcortical structures without much scrutiny.

However, besides the general controversy pertaining to its attentional modulation, the view of an exclusive brainstem localization of the FFR has been recently challenged by Coffey and colleagues (Coffey et al. 2016) for human participants (for similar guinea pig data see (Wallace et al. 2000)). Using MEG and EEG concurrently, they confirmed that the ABR can be acquired with MEG as previously shown by Parkkonen and colleagues (Parkkonen et al. 2009). More importantly, they showed that the FFR can be acquired with MEG as well. Using source projection, FFR activity was present in all auditory subcortical nuclei (i.e. brainstem and thalamic) but most importantly significant auditory cortical contributions were also identified. The latter finding has important implications concerning past studies using the FFR. If the FFR has cortical components, it is certainly possible that any reported attentional effect could have cortical origins instead of or in addition to subcortical ones. It clearly follows that the source of any effect in an FFR paradigm should be determined by source analysis techniques (Hämäläinen and Ilmoniemi 1994; Van Veen et al. 1997; Gross et al. 2001; Lin et al. 2006). Additionally, reporting the onset latency and temporal dynamics of the FFR along with the reported effects becomes crucial, which is not common in current practice (e.g. (Galbraith et al. 2003; Lehmann and Schönwiesner 2014; Varghese et al. 2015)). Studies that report the FFR onset latency estimate it to be between 6 and 10ms (Hoormann et al. 2004; Russo et al. 2004). This time-window overlaps with estimates of the amount of time necessary for the first volley of activity to reach the auditory cortex which is around ~9ms (Liegeois-Chauvel et al. 1991; Brugge et al. 2008, 2009). In order to scrutinize the subcortical and cortical effects of attention on the FFR, we acquired data with whole-head MEG in a cross-modal attention paradigm. We confirmed the strong cortical contributions to the FFR as reported by Coffey and colleagues (Coffey et al. 2016) and furthermore showed that attentional modulations of the FFR affects only cortical regions.

## Materials and Methods

### Participants

38 volunteers (19 females) took part in the experiment and provided written informed consent. At the time of data acquisition, the average age of the participants was 24.4 years (SD: 6.1). Two of these participants had to be excluded because not all six runs were recorded. One participant was excluded because of excessive power in the time frequency data. One further participant was excluded because more than six sensors were marked as bad by the RANSAC algorithm. The final sample of participants included 34 volunteers (19 females) with an average age of 24.4 years (SD: 6.3). All participants reported no previous neurological or psychiatric disorder, and reported normal or corrected-to-normal vision. The experimental protocol was approved by the ethics committee of the University of Salzburg and was carried out in accordance with the Declaration of Helsinki.

### Stimulation Paradigm

Stimulation was controlled by a custom Matlab script using the Psychophysics Toolbox (Brainard 1997; Kleiner et al. 2007). Stimulus presentation and exact timing was ensured by using the VPixx System (DATAPixx2 display driver, PROPixx DLP LED Projector, TOUCHPixx response box by VPixx Technologies, Canada). We used the Blackbox2 Toolkit (The Black Box ToolKit Ltd, Sheffield, UK) to measure and correct for timing inaccuracies between triggers and the visual and auditory stimulation.

The participants performed six runs of a crossmodal attention task (see Figure 1). For each of the 85 trials, an attentional cue indicated whether the participant had to react to a rare oddball in either the visual or auditory domain. Each trial started with a central fixation cross, presented for 500ms followed by the attentional cue (picture of an eye or an ear) presented for 500ms. A fixation cross appeared for 1000ms, followed by the audiovisual stimulation. The auditory stimulation consisted of 30 repetitions of a /da/ sound with an effective fundamental frequency of 114Hz, lasting 40ms (King et al. 2002; Skoe and Kraus 2010). Each presentation of the /da/ sound was followed by 50ms of silence. For the duration of the auditory stimulation, a vertically oriented gabor patch (visual angle: spatial frequency: 0.01 cycles/pixel, sigma: 60) was presented at the center of the screen. 15 of the 85 trials were target trials. If eight target trials had a visual target, the other seven target trials had an auditory target and vice versa. In visual target trials, the gabor patch was tilted by 10° to the left for 270ms anywhere during the presentation time. In auditory trials, three consecutive presentations of the /da/ sound were reversed. The participants had to press a button with their right thumb if the current trial was a target trial of the cued modality. They were allowed to answer as soon as the target occured. After the audiovisual stimulation had finished, participants were given an additional 300ms to answer in order to account for target trials in which the targets appeared towards the end. After each trial, a smiley was presented for 1000ms, indicating whether the (non)response of the participant was correct or incorrect.

**Figure 1:**
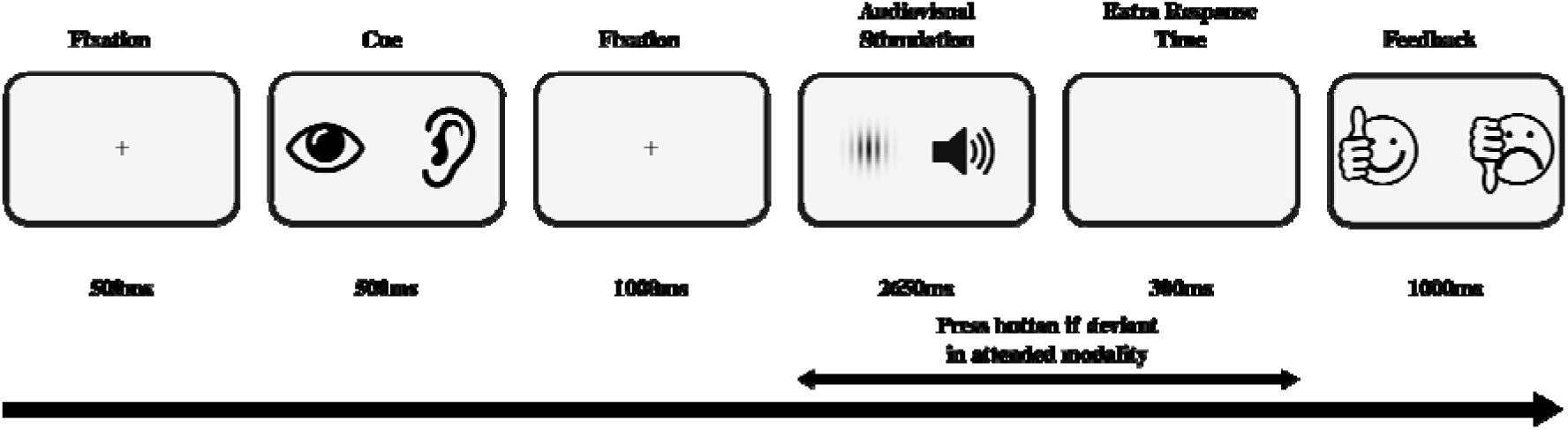
Timeline of the trials of experimental paradigm.

### Data acquisition

Concurrent acquisition of the magnetic and electrical signal was performed at a sampling frequency of 5000Hz (hardware filters: 0.1 - 1600Hz) using a whole-head MEG system (Elekta Neuromag Triux, Elekta Oy, Finland), placed in a magnetically shielded room (AK3b, Vacuumschmelze Hanau, Germany). Brain activity was sampled from 102 magnetometers, 204 orthogonally placed gradiometers and 128 EEG channels. Only the data from the magnetometers are reported due to their greater sensitivity to deep sources as compared to gradiometers. The data quality of the EEG recording was not sufficient for the analysis: movement artifacts were excessive in amplitude, probably due to the participants’ heads touching the surface of the MEG helmet, and many EEG channels showed excessive noise for currently unknown reasons.

### Data preprocessing

Preprocessing of the MEG data was done in a two-step approach. In a first step Signal Space Projection (SSP) was applied to remove exogenous contaminations (Uusitalo and Ilmoniemi 1997). Further data cleaning was performed using a fully automated approach implemented in the autoreject package (version 0.1 running on Python 3.6.8) (Jas et al. 2016, 2017). Specifically, we used autoreject to identify bad sensors and periods containing artifacts, which were subsequently discarded. This approach is detailed in the following paragraph.

Because common artifacts in MEG data are found in rather low frequencies, the data of each run were bandpass filtered between 1-40Hz (FIR filter with hann window, low transition width: 0.1Hz, high transition width: 4Hz, filter length: 165001). The filtered data were then split into epochs of 1s because the algorithms provided by autoreject require epoched data. Each epoch was further downsampled to 500Hz to increase the speed and decrease the computational demands of the artifact identification algorithms. We first applied the RANSAC algorithm to identify sensors that contained data that were highly dissimilar to those of the other sensors (Fischler and Bolles 1981; Bigdely-Shamlo et al. 2015; Jas et al. 2017). If a sensor was marked as bad by the RANSAC algorithm in one run, the sensor was excluded for all runs of the respective participant. If the total number of bad sensors exceeded five, the data of the participant were rejected, leading to the exclusion of one participant. The remaining data were subjected to the “local autoreject” algorithm (Jas et al. 2016, 2017) to identify which of the 1s periods contained artifacts. The exact parameters for the RANSAC and “local autoreject” algorithms can be found in the supplementary material. Each 1s period that was marked as bad by the algorithm was discarded from further analysis. Subsequent analysis on cleaned data was carried out using the open source MNE-Python toolbox (version 0.17.2 running on Python 3.6.8) (Gramfort et al. 2013, 2014). Statistical analysis was performed using Eelbrain version 0.29.5 (Brodbeck n.d.). Perceptually uniform colormaps ((Kovesi 2015); (Bedna n.d.)) were used for all color-coded figures.

We bandpass filtered (FIR filter with hann window, passband: 80-2000Hz, low transition width: 5Hz, high transition width: 100Hz, filter length: 3301) the raw data, followed by a bandstop filter to eliminate line noise contamination (FIR filter with hann window, stop bands at 50Hz and multiples up to 1950Hz, transition width: 1Hz, filter length: 33001). We extracted epochs of data in the time window of 60ms before to 120ms after the onset of each individual auditory stimulus, accounting for a 16ms delay introduced by the tubes of the MEG-proof sound system and 7ms delay inherent to the sound file we used. In order to reduce possible contamination of the auditory signal by the visual evoked response, the first four auditory stimuli of every trial were discarded. We further discarded all target trials and trials in which a false positive response was given by the participant. The remaining trials were averaged within their respective condition (attend visual / attend auditory) in order to compute the attention effect. A further average of all trials of each participant was calculated in order to locate the FFR in time, frequency and space.

### Sensor Space Analysis

We applied a wavelet transform around the fundamental frequency of the stimulus (Morlet Wavelets, six cycles, 104-124Hz, Δf=2Hz, Δt=1ms) on the averaged data. In order to exclude any remaining outliers from the analysis, the resulting power values were averaged within each participant. The individual power values were Z-transformed and those participants whose average power was three standard deviations below or above the group average were excluded. This lead to the exclusion of one participant (z=4.89). We computed the power envelope of the FFR by first averaging over all frequency bins.

In order to visualize the FFR on sensor level, the power values were first averaged over all remaining magnetometers within every participant. The average power and standard deviation over all participants was subsequently calculated and resulted in an FFR response peaking at 51ms after stimulus onset as shown in Figure 2a.

**Figure 2:**
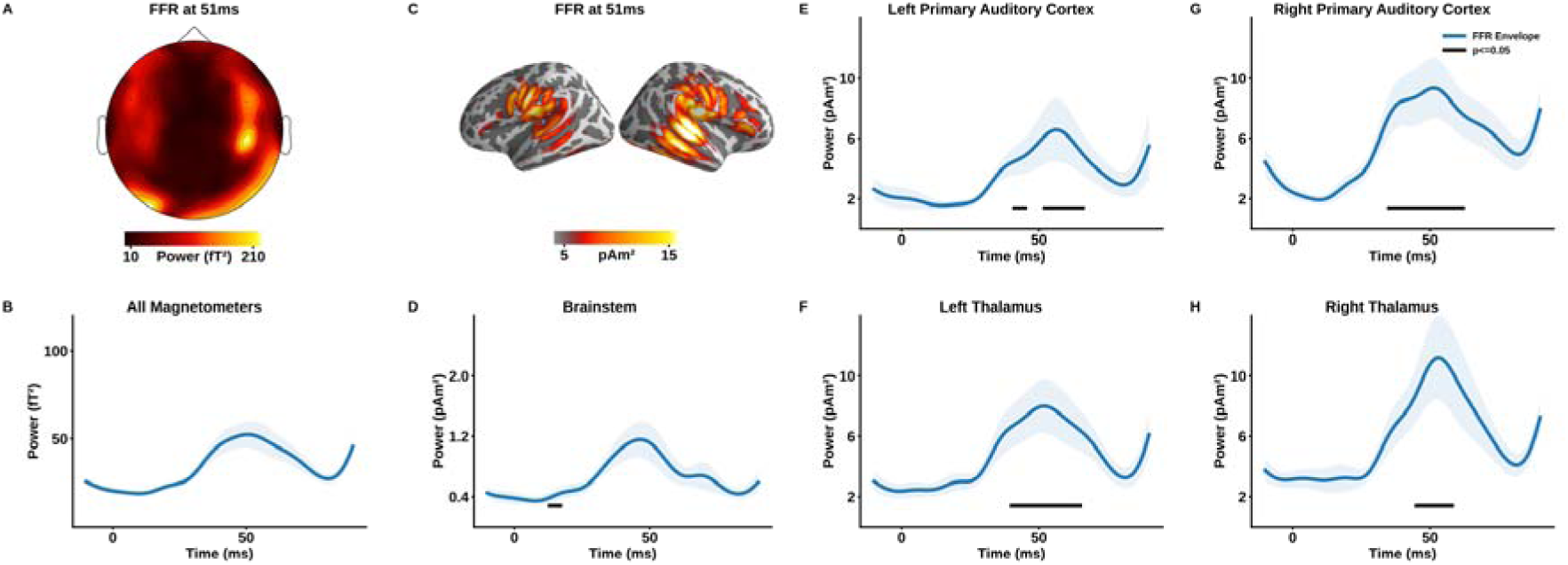
The Frequency Following Response (FFR) in sensor and source space. A) Topography of the FFR at the time of maximum power. B) Timecourse of the evoked response at the fundamental frequency (f0) of the stimulus averaged over all magnetometers. Shaded error bars denote the standard error. C) Source reconstruction of the FFR at the time of maximum power. D-H) Timecourse of the evoked response at the f0 for the 5 ROIs. Shaded error bars represent the standard error. The black bar at the bottom of each panel represents the temporal extents of the clusters in which the respective ROI showed a higher response than the orbitofrontal cortex, used as a control region.

In order to test our principle hypothesis that FFR activity was higher when the auditory modality was attended, we applied a cluster-based nonparametric, threshold free permutation-based statistic (Maris and Oostenveld 2007; Smith and Nichols 2009) (dependent samples t-test, 10000 permutations, channel neighborhood structure provided by MNE Python) to the data, restricted to the time-window between stimulus onset to 90ms later.

### Source Space Analysis

Nine of the 35 participants included in the final analysis provided us with high-quality T1 MR images. These MRIs were segmented with Freesurfer (Fischl 2012). Alternatively, the average brain provided by Freesurfer (fsaverage) was morphed to the participants’ headshape. The surface of the inner skull was either extracted using Freesurfer (Fischl 2012) if the individual anatomical images were available or determined by applying the transforms used for morphing the average brain to the participants’ headshapes. The coordinate frames of the MR images and the MEG sensor positions were coregistered using MNE Python (Gramfort et al. 2013, 2014). We subsequently computed a one-layer boundary-element model (BEM) (Akalin-Acar and Gençer 2004) to accurately model the propagation of magnetic fields from generators in the brain to the sensors. We constructed the cortical source space using 4098 sources, each covering approximately 24mm^2^. In order to estimate activity at the brainstem and the thalamus, the surface-based source space was combined with a volumetric source space, placing equidistant source of 5mm spacing in the regions labeled “Brainstem”, “Left-Thalamus-Proper” and “Right-Thalamus-Proper”, as defined in the “aseg” atlas (Filipek et al. 1994; Seidman et al. 1999; Fischl 2012). Further cortical regions of interest (ROI) were defined using the HCP-MMP1.0 atlas (Glasser et al. 2016) morphed to the individuals’ anatomy. Similar to Coffey and colleagues (Coffey et al. 2016), we used the Primary auditory cortex (A1) as the cortical ROI and defined the orbitofrontal cortex (OFC) as the control region. Evoked sensor space activity was projected to the defined sources using the Minimum Norm Estimate method (Hämäläinen and Ilmoniemi 1994) with a depth weighting coefficient of 0.8 (Lin et al. 2006). We subsequently applied a wavelet transform with the same parameters used in the sensor space analysis to all orientations of every source the data were projected to. The power values within each source were combined by summing the values of the three orientations. As for the sensor space data, we obtained the power envelope of the FFR by averaging the power over all frequency bins. This approach resulted in data for sources on the cortical surface and the previously defined subcortical regions. In order to assess cortical contributions to the FFR and the attention effect, we morphed these cortical sources to the average brain (fsaverage) provided by Freesurfer. To visualize the cortical attention effect, we calculated T-values (dependent samples, one tailed). For the subsequent ROI analysis, we averaged the power time-courses of each source belonging to each of the ROIs defined above.

The first question we wanted to answer was whether FFR-related activity was higher in the cortical and subcortical auditory regions compared with the two control regions (OFC, left and right hemisphere). We therefore averaged the power of the two control regions. We then applied the same cluster-based, threshold-free permutation statistics (Maris and Oostenveld 2007; Smith and Nichols 2009) (dependent samples t-test, 10000 permutations, one-tailed) that we used for the sensor data to contrast the power of each of the putatively auditory ROIs with the power of the averaged control region.

The second and crucial question we tried to answer was which of the ROIs were affected by the attentional modulation. We therefore contrasted the power during auditory vs. visual attention at each of the ROIs individually, again using the same cluster-based, threshold-free permutation statistics (Maris and Oostenveld 2007; Smith and Nichols 2009).

## Results

### Behavioral Results

Behavioral response data showed that participants gave the correct response in 99% (SD: 0.7%) of the trials. When a target was presented in the cued sensory domain, participants correctly gave a response in 95% (SD: 3.1%) of the respective trials. False responses to targets without triggers were given in only 0.1% (SD: 0.3%) of the trials.

### Sensor space analysis

In a first step, we analyzed the temporal dynamic of the FFR in sensor space and subsequently compared the data acquired during the “attend auditory” and the “attend visual” condition. The power envelope showed an evoked response to the sound stimulus peaking at ~51ms after stimulus onset (see Figure 2B). The topography of the response shows a bilateral activation pattern, mostly over temporal regions, lateralized to the right hemisphere (see Figure 2A). Since meaningful control regions cannot be defined on the sensor level, we refrained from further statistical analysis.

The cluster-based nonparametric, threshold-free permutation-based statistics for the impact of the attentional modulation shows that the FFR response is significantly larger when attention is focused on the auditory domain (p=0.035, see Figure 3B). The effect is, however, restricted to a rather short and late period after stimulus onset (66ms - 74ms). It also does not coincide with the maximum of the FFR itself. This low power and specificity is likely due to an interaction between the low spatial specificity of the magnetometers and the fact that the average over all magnetometers was analyzed. Figure 3A shows the topography of the comparison between the two conditions at the time of the maximum power of the FFR. It suggests a weak overlap with the topography of the FFR itself (see Figure 2A). This analysis confirms the presence of the FFR in sensor space and is suggestive yet not conclusive with respect to its modulation by selective (intermodal) attention.

**Figure 3:**
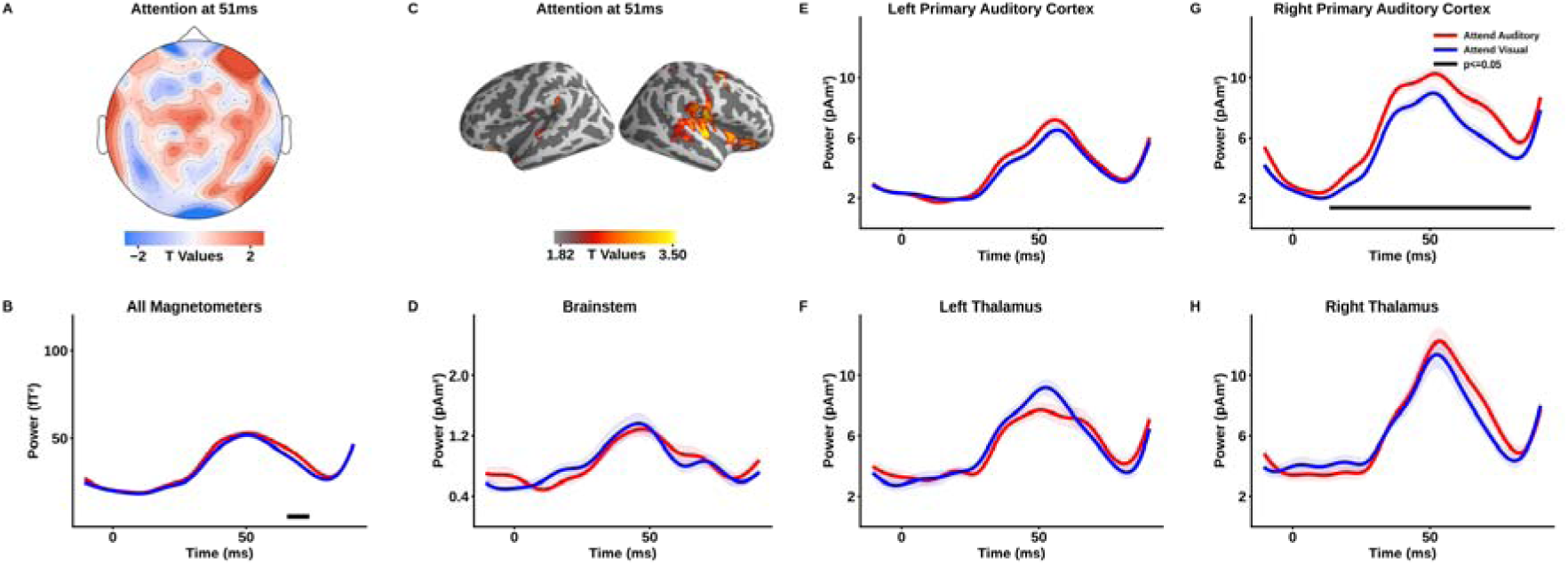
The attention effect in sensor and source space. A) Topography of the attention effect at the time of maximum power of the FFR. B) Timecourses of the FFR when attention was directed to the auditory domain and the visual domain. Shaded error bars denote the standard error of the mean for within-subjects designs (Morey and Others 2008). The black bar at the bottom of the panel represents the timeframe in which the FFR was higher when the auditory domain was attended. C) Source reconstruction of the attention effect. D-H) Timecourse of the FFRs to both conditions. For details, refer to the description of panel B.

### Source space analysis

After confirming the presence of the FFR as well as finding suggestive evidence for the hypothesized attentional modulation in sensor space, the next step was to locate the respective generators. Importantly, we wanted to reveal possibly different contributions by cortical and subcortical sources. Source projection of the FFR showed strong cortical contributions, lateralized to the right hemisphere (See Figure 2C). The maximum power was found at the Auditory 4 complex, an auditory region ventral to the primary auditory cortex. The activity is rather widespread, including the primary auditory cortex as well as ventral temporal, parietal and prefrontal regions. The ROI analysis statistically confirmed this notion (see Figure 2D-H). All five ROIs showed responses to the stimulus at its fundamental frequency. In order to quantify whether the response was specific to the ROI, the envelope at each ROI was statistically contrasted to the envelope at a control region in the orbitofrontal cortex. This analysis showed that all ROIs generated a significantly stronger FFR than the control region (see Figure 2). The time periods of the significant increases of all ROIs except the Brainstem were well within the range of the sensor-level FFR. The Brainstem ROI on the other hand showed the familiar peak at approximately the same time as the sensor level data and the other four ROIs. However, significant FFR increases were obtained only for a period between 13ms and 18ms after stimulus onset (for details see Supplementary Table 1).

The cortical areas showing attentional modulation of the FFR are mostly restricted to the right hemisphere and cover early auditory regions as well as regions in the temporo-parieto-occipital junction, the lateral prefrontal cortex and ventral parietal areas (see Figure 3C). These regions also strongly overlap with those found to show general FFR-related activity. The ROI analysis indicated that the attention effect was exclusive to the right primary auditory cortex (p=0.0005). Although Figure 3 descriptively suggests a trend for the right thalamus, the attention effect fails to reach significance at that area (p=0.203). This is also the case for the left thalamus (p=0.619), the brainstem (p=0.402) and the left primary auditory cortex (p=0.164).

These results strongly indicate that although we were able to record subcortical — especially thalamic — contributions to the FFR, only cortical contributions to the attentional process could be found in the data.

## Discussion

It is commonly accepted that the FFR can be used as a proxy to subcortical activity which is otherwise hard to detect in the MEG and EEG (Chandrasekaran and Kraus 2010; Skoe and Kraus 2010). The rationale behind this assumption is that only subcortical areas exhibit the strict phase-locking behavior at frequencies above 100Hz (corresponding to the typical F0) that are commonly used for the stimuli and that the onset latency is too early for cortical generators (Wallace et al. 2000; Skoe and Kraus 2010). These assumptions have been recently challenged by a study by Coffey and colleagues (2016) that shows strong cortical contributions to the FFR and by studies showing that the auditory cortex reacts to sound stimulation as early as 8-9ms after stimulus onset (Liegeois-Chauvel et al. 1991; Brugge et al. 2008, 2009). The quasi-automatic attribution of experimental effects on properties of the FFR to subcortical areas thus needs to be revisited.

The FFR has been widely used to study top-down effects of attention on subcortical auditory areas with inconsistent results (Galbraith et al. 2003; Hoormann et al. 2004; Lehmann and Schönwiesner 2014; Varghese et al. 2015; Forte et al. 2017). Yet, none of these studies used current source projection algorithms to estimate the location of the generators of the reported signals and only some (Hoormann et al. 2004; Russo et al. 2004; Forte et al. 2017) showed the temporal dynamics. The current study is the first that uses the unique strength of the MEG — with its excellent temporal resolution and good spatial resolution — to scrutinize the spatial location of a possible attention-related effect on the FFR without assuming the location of its generators a priori. In sensor space, the attention effect is found considerably later than the FFR, already pointing towards cortical generators. In fact, the simple FFR activation as well as the attention effects outlast the actual period of stimulation, arguing for some reverberatory processes. Overall, the source space analysis confirms the sensor-level finding but also significantly expands it. Firstly, we show next to subcortical generators (brainstem at early and thalamus at later time-periods) of the FFR also strong contributions of auditory cortex. This part of the study confirms that the FFR can be detected using MEG (Coffey et al. 2016) as well as recent findings that the origin of the FFR is not restricted to subcortical areas (Wallace et al. 2000; Coffey et al. 2016). Also the temporal evolution of the FFR in line with the study by Coffey and colleagues, with a peak reached at ~51ms, with the exception of brainstem where significant activation (as compared to the control region) was identified significantly earlier at ~13ms. These results may point to indeed an earlier “feedforward” projection of phase-locked activity involving the brainstem, whereas the later portions of the FFR (including those following the stimulus offset) may be more driven by cortico-thalamic interactions. These aspects are beyond the scope of the current study, but open up interesting perspectives in future research on the FFR. Most importantly, however, apart from largely confirming the FFR generating structures as suggested by Coffey and colleagues, we show that only the right primary auditory cortex shows a significant effect of attention.

Of course, our results do not disprove the presence of attention-related effects on subcortical regions in general and on subcortical generators of the FFR in particular. Sufficient evidence for attention effects in auditory subcortical areas and even the cochlea is available (Giard et al. 1994; Maison et al. 2001; Raizada and Poldrack 2007; Rinne et al. 2008; Wittekindt et al. 2014; Slee and David 2015; Riecke et al. 2018). The perfectly regular stimulus presentation at 11.1Hz might have lead to cortical entrainment, boosting the signal-to-noise ratio of the cortical generators. However, as stated in the introduction, existing reports on the presence or absence of attentional effects on the FFR used EEG setups that strongly assumed the absence of cortical generators to the FFR (Galbraith et al. 2003; Hoormann et al. 2004; Gutschalk et al. 2008; Lehmann and Schönwiesner 2014; Varghese et al. 2015). The most prominent “optimization” of the recording setup was that only very few electrodes were used. If the assumption of the absence or irrelevance of cortical generators were true, the number and the location of the electrodes should not be relevant. If, as shown by our results, the FFR evokes strong cortical activity and attention-related effects are strong in auditory cortical regions, the small number of electrodes and their possibly inconsistent locations would be highly relevant and could, at least in part, explain the inconsistent results.

To conclude, by recording the FFR with the MEG at high temporal and spatial resolution during a cross-modal attention task and using state-of-the-art source projection techniques, we confirm that the cortical generators of the FFR exist and demonstrate for the first time that they are modulated by attention. The lack of an a-priori assumption on the generators of the FFR allowed us to provide a more differentiated perspective on the underlying sources of the FFR and its role in auditory processing. Our results strongly suggest that high-density recording and source projection techniques should be used in future research to disentangle the diverse contributions from cortical and subcortical regions.

## Acknowledgements

The authors wish to thank David Opferkuch and Manfred Seifter for their assistance during the collection of the data. We would like to also thank Hayley Prins for proofreading the manuscript. Address of corresponding author: Dr. Thomas Hartmann, Department of Psychology, University of Salzburg, Hellbrunnerstr. 34, 5020 Salzburg, Austria.

## Supplementary Table

**Supplementary Table 1:** Time Periods of significant FFR for each ROI

| ROI | Significant Time Periods | p-Values |
| --- | --- | --- |
| Left Auditory Cortex | 41ms - 46ms | 0.010 |
|  | 52ms - 67ms | 0.024 |
| Right Auditory Cortex | 35ms - 63ms | 0.007 |
| Left Thalamus | 40ms - 66ms | 0.003 |
| Right Thalamus | 45ms - 59ms | 0.013 |
| Brainstem | 13ms - 18ms | 0.037 |

### Parameters of autoreject

RANSAC: n_resample=50, min_channels=0.25, min_corr=0.4, unbroken_time=0.4 Local autoreject: n_interpolate=[1, 4, 32], consensus=[0, 0.25, 0.5, 0.75, 1]

